# Embryonated chicken eggs clear systemic H3N2 influenza without RIG-I: transcriptomic evidence for innate sufficiency and brain immune privilege

**DOI:** 10.64898/2026.05.11.723864

**Authors:** Ozlem Gurses, Janice B Fung, Deepali Lehri, Ravi Sachidanandam

## Abstract

An apparent paradox drives this study: H3N2 influenza virus concentrates in the brain of infected 10-day chicken embryos while kidney and lung, which express the same viral entry receptors (ST3GAL3 and other sialic acid receptors), are essentially virus-free. Using mRNA-seq on brain, kidney, and lung from H3N2-infected 10-day chicken embryos, we establish and propose a resolution to this paradox, by identifying immune privilege, rather than neurotropism, as the more likely explanation; circulating macrophages likely clear the virus from peripheral tissues but seemingly cannot cross the embryonic brain barrier. The innate response is robust despite lacking RIG-I: MDA5/IFIH1 and TLR3-TLR7-IRF signaling compensate robustly, likely driving complete viral clearance in peripheral organs. At 48 h post-infection, macrophages in kidney and lung are in a post-clearance M2 state; complement is activated but lacks both the H3N2-specific antibodies and the terminal C9 component for productive effect. These findings challenge the hypothesis that RIG-I loss necessarily renders chickens more susceptible to influenza: complete peripheral clearance of a productively infecting H3N2 strain in the absence of RIG-I is difficult to reconcile with an obligate role for RIG-I in this host’s antiviral defense. We also identify the embryonic brain as a potentially immune-privileged viral sanctuary with implications for influenza neurological disease in young hosts.

## 2 Introduction

Chickens are central to global food security and serve as reservoir hosts at the avian-human influenza interface[1]. Their embryonic immune system develops along a well-defined timeline: interferon (IFN) responses emerge by days 7-10, T cells by day 11, B cells by day 12, with full adaptive competence approximated by day 18[2, 3]. Before B-cell maturation, maternal IgY transferred from the yolk initiates humoral coverage. This staged development makes early embryos ideal for isolating innate from adaptive contributions to antiviral defense.

Innate antiviral immunity in mammals is mediated by pattern recognition receptors (PRRs) including TLRs, RIG-I-like receptors (RLRs), and the cyclic GMP-AMP synthase-stimulator of interferon genes (cGAS-STING) axis[4]. A recurring hypothesis in the field holds that Galliformes are intrinsically more susceptible to influenza because they lack one of these sensors, RIG-I (DDX58) which is a cytosolic sensor that detects viral RNA and initiates type I interferon production in mammals[5, 6, 7]. This framing implies a fundamental innate sensing gap that could explain why chickens serve as amplifying hosts at the avian-human interface[8, 9, 10, 11].

The TLR-IRF axis is a central component of this compensatory architecture. Ten chicken Toll-like receptors (TLRs) mediate innate sensing, including the avian-specific TLR15 and TLR21[12], the latter functionally replacing mammalian TLR9 for nucleic acid detection. TLR3 and TLR7 sense dsRNA and ssRNA respectively, and signal through IRF1 and IRF7 to induce type I IFNs and activate NF-*κ*B[13]. In the cytosol, IFIH1 (MDA5), a paralog of RIG-I sharing its RNA-sensing helicase domains, likely assumes RIG-I’s role in detecting viral RNA and initiating downstream signaling through MAVS[5, 6]. IRF7, a key transcription factor for early innate responses in chicken macrophages, drives upregulation of ISGs (PKR, OAS, viperin) and IFITMs that halt viral replication[13, 14], providing a compensatory sensing axis in the absence of RIG-I.

Prior work characterized antiviral responses to Newcastle disease virus (NDV) and avian influenza using targeted assays[2, 3, 15]. To our knowledge, this is one of the first studies to apply unbiased mRNA-seq simultaneously across multiple tissues of the embryonated egg, using H3N2 (A/Uruguay/716/2007) which is the subtype responsible for the 1968 pandemic and a continuing cause of seasonal influenza with documented capacity to infect avian hosts[16].

At day 10, functional T and B cells have yet to develop, isolating the innate response, while the brain and metanephric kidneys are undergoing active organogenesis[17, 18], making this stage ideal for simultaneously studying antiviral immunity and its developmental context. Profiling expression in brain, kidney, and lung in parallel allows us to distinguish local from systemic responses and, critically, to identify which tissues harbor infecting virus and which have cleared it. By integrating differential expression and pathway enrichment across all three tissues, we define shared and tissue-specific response modules, catalog the functional compensators for absent mammalian immune genes (Table ST2), and provide evidence that the embryonic brain is immune-privileged in a way that permits unchecked viral infection when systemic clearance is otherwise complete.

Our data directly challenge the idea that the absence of RIG-I introduces a vulnerability; viral clearance in peripheral tissues is complete despite RIG-I absence, and the downstream ISG response is fully activated, demonstrating that the avian innate immune system has seemingly evolved a parallel and equally effective sensing architecture. This is further supported by previously published experiments showing that reinstating RIG-I in chickens paradoxically increases inflammation[19], and by suggestions that RIG-I loss might confer selective advantage by modulating interferon-driven inflammation[20].

## 3 Materials and Methods

### 3.1 Embryonated Eggs

Ten-day-old specific pathogen-free (SPF) embryonated chicken eggs (ECEs; Pocono Rabbit Farm & Laboratory, Canadensis, PA) were used under institutional guidelines approved by the Institutional Animal Care and Use Committee (IACUC) of New York Medical College[21]. Embryonated eggs were obtained as surplus material from an ongoing study conducted under that approval; no additional animals were procured or sacrificed for the present study.

### 3.2 Influenza Virus

We used a reassortant H3N2 variant, NYMC X-175C, bearing the hemagglutinin (HA) and neuraminidase from A/Uruguay/716/2007 and the M protein from A/Puerto Rico/8/1934, the latter included for improved growth [22]. This strain carries the well-characterized L194P substitution in HA protein, which shifts receptor-binding specificity toward the avian-type *α*2,3-linked sialic acids present in chicken tissue [23, 24], enabling productive replication in embryonated eggs. We selected this virus for its demonstrated capacity to infect embryonated eggs and because it is already used at our institution for vaccine seed-virus production, allowing us to draw on established infrastructure and expertise for this system.

#### Plaque assay to establish stock concentration

MDCK-SIAT1(ATCC CRL-3743) monolayers were infected with 400 µL of serially diluted virus suspension for 60 min at 37°C, 5% CO_2_, with intermittent rocking to allow adsorption. Inoculum was then removed and monolayers overlaid with 2% agarose (SeaKem LE, Lonza Bioscience, Morrisville, NC) in autoclaved deionized H_2_O containing TPCK-treated trypsin, which enables cell-to-cell spread of progeny virus while restricting diffusion so that plaques remain spatially discrete; 8 mL of overlay was added per dish. Dishes were incubated at 37°C, 5% CO_2_ for 72 h, after which agarose plugs were removed and monolayers stained with 0.1% crystal violet in 20% ethanol. Infected cells lyse and fail to stain, appearing as clear plaques against the stained monolayer; each plaque is assumed to derive from a single infectious virion. Titer was calculated as PFU/mL = *N/*(*V xD*), where *N* is the mean plaque count across three technical replicates, *D* is the dilution factor, and *V* is the inoculum volume (in mL). Our strain, A/Uruguay/716/2007 (H3N2), yielded 60-65 countable plaques at a 10*^−^*^5^ dilution, corresponding to 1.5*x*10^7^ PFU/mL (Figure SF1).

#### EID50

Viral dosage was determined as follows. Starting viral titer was established by plaque assay on MDCK-SIAT1 described above (Figure SF1). Serial 10-fold dilutions (10*^−^*^1^ *−* 10*^−^*^7^) of virus, alongside mock, were inoculated into the allantoic cavity of groups of twelve 10-day-old SPF embryonated eggs, sealed with wax [25], and incubated for 48 h at 37°C [26, 27]. Allantoic fluid was harvested and stored at 4°C. The 50% egg infectious dose (EID_50_), the dilution producing detectable HA activity in 50% of inoculated eggs (6/12), was 10*^−^*^5^, corresponding to a titer of 10^5^ EID_50_/mL[28, 29]. Infection was independently confirmed by RT-PCR for viral HA gene and PB1 transcripts (Figure SF1).

#### Hemagglutination assay (HA assay)

Serial two-fold dilutions of virus suspension in PBS were mixed with chicken erythrocytes to determine the viral titer, the reciprocal of the highest dilution producing visible agglutination, as a measure of receptor-binding activity of the viral preparation.

### 3.3 Infection protocol

Mock infection was carried out with 100 µL of PBS containing 25µg/mL gentamicin. For viral infection, based on the EID_50_ determined above, 100µL of PBS containing 150 PFU/mL of H3N2 (NYMC X-175C), corresponding to *≈* 15 PFU per egg, was used (Figure 1). HA assays were used to determine viral activity and concentration. The HA assay works as follows; Hemagglutinin on the surface of viral particles binds to sialic acid receptors on chicken red blood cells (RBC) which have been isolated, washed and purified from blood. Without the binding to hemagglutinin, the RBC clump together and sediment as a red dot at the bottom of the well (signalling a lack of sufficient virus in the sample), but the binding to hemagglutinin of the virus keeps the RBC in suspension in a diffuse matrix. By creating a serial dilution, and testing against the RBC, the highest dilution that prevents clumping (appearance of red dot at bottom of well), measures the concentration of virus in the sample.

**Figure 1:**
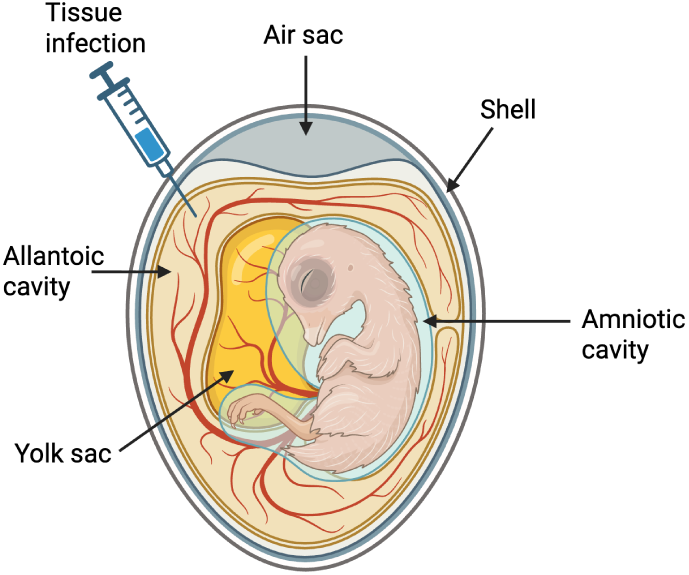
Experimental design. Schematic of an embryonated chicken egg. Eggs used in our study were raised to be Specific Pathogen Free (SPF). At 10 days old, the allantoic fluid was infected with *≈* 15 PFU (100 µL of a 150 PFU/mL preparation, corresponding to the EID_50_ dilution of the stock) to cause infection without being immediately fatal to the embryos. Control eggs were injected with PBS containing 25µg/mL gentamicin. Hemagglutination and plaque assays confirmed viral titer and viability of the inoculum; RT-qPCR of allantoic fluid confirmed productive in ovo replication following inoculation (Figure SF1). We recover viral transcripts by RNA-seq, in a differential manner across tissues, which is only possible if the virus is actively infecting rather than merely present as residual inoculum in the allantoic fluid.

**Figure 2:**
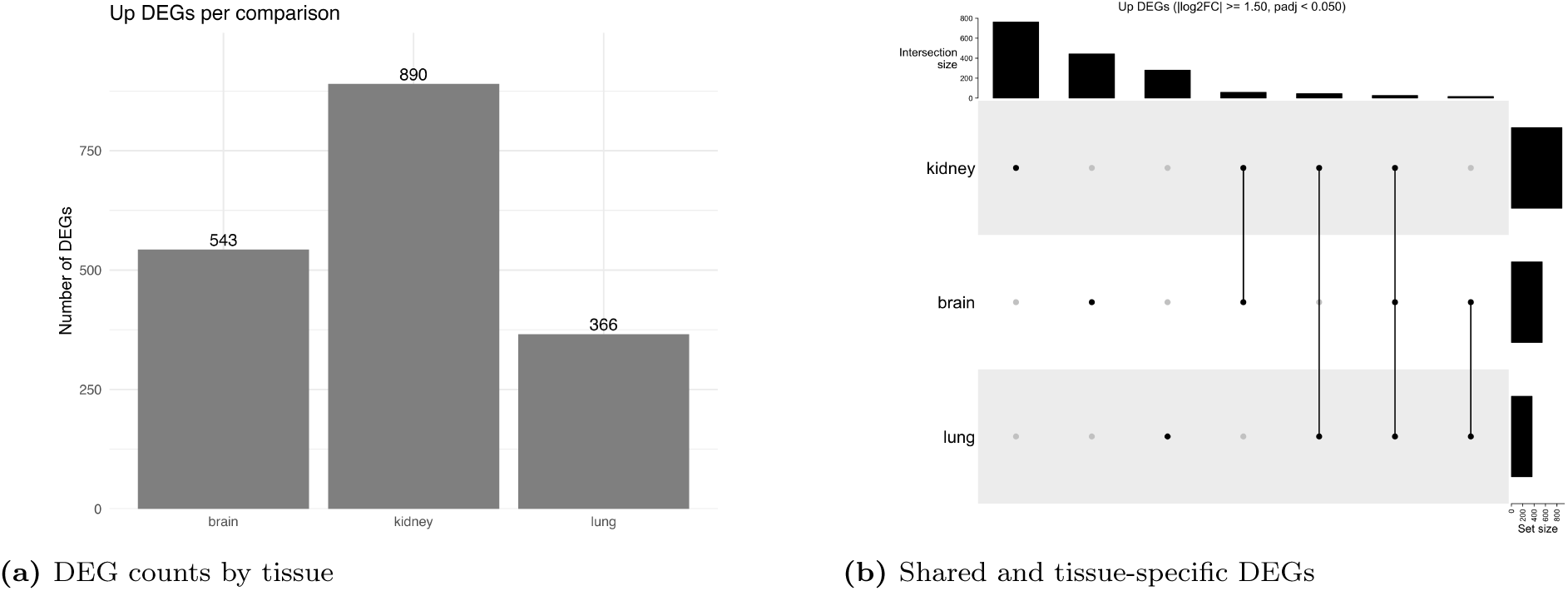
Transcriptome-wide response to H3N2 infection across three tissues. **a)** Number of upregulated differentially expressed genes (DEGs; adjusted log_2_ FC *≥* 1.5) per tissue in infected vs. uninfected embryos. The brain and kidney have many genes that are upregulated under infection; the lung, as we discuss in Figure 3, is not consistently dissected, so is a mix of tissues leading to gene expression varying considerably between them, so a lower number of genes are consistently upregulated in the infected samples, many are associated with common elements like ISGs or macrophage genes which are present across tissues. **b)** UpSet plot of shared and tissue-specific upregulated DEGs across brain, kidney, and lung, illustrating the extent of systemic vs. local transcriptional coordination.

HA assays were performed on allantoic fluid from infected and mock-infected eggs to confirm viral activity prior to tissue harvesting[30]. Additional confirmation of viral titer and in ovo replication were carried out by plaque-forming assays on MDCK-SIAT1 cell monolayers and RT-PCR quantification of HA and PB1 mRNA in allantoic fluid, including comparison of anti-M1 antibody-treated versus isotype-control-treated samples to confirm replication-dependence of the signal(FigureSF1).

### 3.4 Tissue Collection and RNA extraction

Brain(2 control and 2 infected), kidney(2 control, 3 infected), and lung (2 control, 2 infected) tissues were dissected from embryos following chilling at 4*^◦^*C. Total RNA was extracted using TRIzol reagent (1 mL per 50–100 mg tissue) according to standard protocols[31].

### 3.5 mRNA Sequencing

RNA quality and quantity were assessed prior to library preparation. mRNA-seq libraries were prepared using the SMART-Seq mRNA LP kit (Takara Bio; Cat. 634762/5/6), incorporating oligo-dT capture, full-length cDNA synthesis, fragmentation, and indexed adapter ligation. Libraries were sequenced in paired-end mode on an Illumina NextSeq 2000 platform.

### 3.6 Analysis of mRNAseq data

Reference sequences for *Gallus gallus* transcripts were compiled from multiple sources, including the UCSC Genome Browser, NCBI GenBank, and published datasets, and consolidated into a non-redundant reference transcriptome to maximize annotated gene coverage. This was a semi-manual process involving remapping unidentified reads to the genome, and using the underlying annotations (gene predictions, EST sequences etc.) to extract sequences for our mRNA database. We also added the sequences from the H3N2 virus genome used in our experiments, to the database, in order to aid the identification of viral expression.

Gene names in chicken were normalized by a committee in 2009[32]. Since then a number of chicken/galleoform-specific genes have been identified, such as SCARI94 which was called PIT54 for a while, and plays the role of CD163 in mammals. The naming/identity of genes is somewhat in flux. Gene names were assigned to sequences in our transcript database based on either nomenclature used in the Refseq database, or else through homology. The mapping of genes to relevant pathways were done using functional annotations in mammalian systems, as well as manual curation, like in the case of the Scavenger Receptor (SCAR) family of genes which are important for our system, as they are critical for innate immunity, pathogen clearance, and tissue homeostasis.

#### 3.6.1 RNA-Seq Quantification and Differential Expression Analysis

Transcript-level abundance was quantified from single/paired-end sequencing reads against the reference mRNA transcriptome using kallisto (v0.50) [33]. Kallisto reports normalized TPM values for each transcript. Transcript abundances and effective lengths were summarized to gene-level count matrices using the tximport package (v1.30+) [34] in R (v4.3) [35], using a custom transcript-to-gene mapping table extracted from FASTA transcript headers.

A cautionary note: because TPM is a compositional measure, and brain, kidney, and lung differ substantially in transcriptome composition, exact fold-differences in viral TPM between different tissues should not be over-interpreted. Comparisons within a given tissue, such as infected versus its own matched uninfected control, are not subject to this concern, since the composition of a given tissue type is similar across control and infected samples. Our claims of tissue-specific viral burden (Figure 4, Section 4.4) rely on this within-tissue infected-vs-control contrast rather than on direct cross-tissue TPM magnitude comparisons.

Low-count genes were filtered prior to downstream testing by retaining features with *≥* 10 counts in at least two samples. Differential gene expression analysis was conducted using the negative binomial generalized linear model framework implemented in DESeq2 (v1.42) [36]. To evaluate baseline and infection-induced transcriptomic shifts across brain, kidney, and lung tissues, both a factorial interaction model (*∼* tissue + condition + tissue : condition) and a unified group factor model (*∼* group) were constructed. Dispersion estimates and log_2_ fold changes (log_2_ FC) were calculated using empirical Bayes shrinkage where appropriate.

For exploratory data visualization, count data were normalized via variance-stabilizing transformation (vst). Sample relationships and clustering behavior were assessed through Principal Component Analysis (PCA) and sample-to-sample Euclidean distance heatmaps. Differentially expressed genes (DEGs) for each tissue were defined as those exhibiting a Benjamini–Hochberg adjusted *p*-value (*p*_adj_ *<* 0.05) and *|* log_2_ FC*| ≥* 1.5. Overlapping and tissue-specific upregulated gene sets were analyzed and visualized using UpSetR [37]. Expression heatmaps displaying row-wise *Z*-score normalized values were generated using pheatmap [38], applying correlation distance and Ward’s hierarchical clustering (ward.D2). Differentially expressed genes (DEGs) were determined and used to delineate pathways that are activated in different tissues in response to the infection. Viral sequences were identified in infected samples.

## 4 Results

mRNA-seq profiling revealed a coherent picture with five intersecting themes: (1) viral distribution confined to the brain, interpreted as reflecting immune privilege rather than neurotropism; (2) robust antiviral sensing through MDA5/IFIH1 and TLR–IRF signaling despite the absence of RIG-I, driving complete viral clearance in peripheral organs; (3) a post-clearance restorative program dominated by M2-polarized macrophages, iron sequestration and heme scavenging, and a complement activation that is ineffective in this context (lacking H3N2-specific maternal antibodies); (4) A lack of microglial response in the infected brain; and (5) preferential autophagy over apoptosis as an embryo-protective cell-fate strategy throughout.

### 4.1 Tissue dissection evaluated by tissue-specific expression

We identify tissue-specific transcripts that are consistently high across all samples of a tissue (controls and infected) to distinguish the brain (PAX6, SIX3, STMN3) and kidney (SULT,FGG,FGB) samples. This consistency suggests that the dissection of these tissues comprise roughly the same tissue or mix of cells and gives us confidence in the pathway analysis derived from the data for those samples (Figure 3).

**Figure 3:**
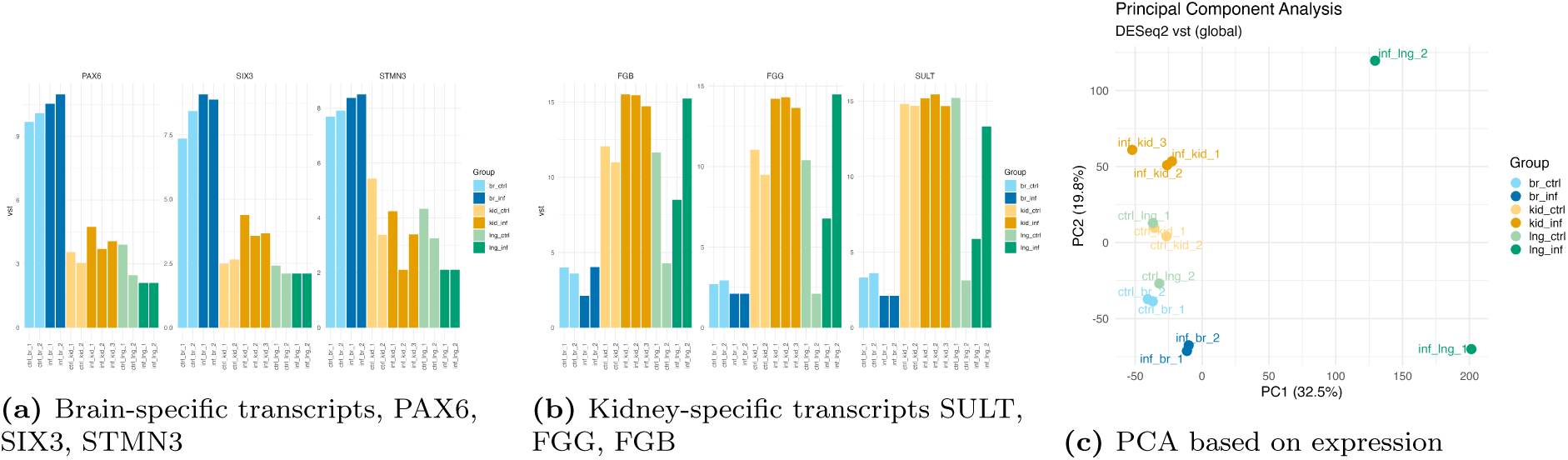
Tissue-specific expression to confirm dissection of tissues. **a) Brain-specific genes** PAX6, SIX3, and STMN3, are highly expressed in infected and control samples, suggesting that these have sampled a reasonably similar set of cells that are derived from brain tissue. **b) Kidney-specific genes** SULT, FGG, and FGB, are highly expressed in infected and control samples, suggesting that these have sampled a reasonably similar set of cells that are derived from kidney tissue. **c) PCA of samples** The brain and kidney samples cluster as expected, by tissue and sub-cluster by infection status. But the lung samples are widely separated, suggesting they are not a pure sample but a mix of cell types. Not many lung-specific transcripts are dominant in the lung isolates, the lung is not well-formed and our isolates are likely a mix of tissue types.

In contrast, the lung samples showed a lot more variability, leading us to be more cautious in interpreting the lung samples. Lung dissection at this embryonic stage lacks precise anatomical landmarks, potentially introducing sampling variability into lung gene expression data. The data from PCA cluster (Figure 3) is in agreement with this interpretation. This leads us to conclude they likely represent a mixture of respiratory and adjacent tissue; So we treat ”lung” as a proxy for peripheral, non-CNS tissue rather than a molecularly pure lung sample (Figure 3).

### 4.2 Confirmation of infection

Sequencing data from mRNAseq has evidence of viral transcripts in the infected samples which suggests active infection of the embryo. Additional confirmation is provided by the HA assay, antibody staining and RT-PCR (FigureSF1).

### 4.3 Overview of Transcriptional Responses

Differential expression analysis identified substantial and tissue-specific transcriptional responses across all three tissues (Figure 2). The brain and kidney (more so) had a large number of upregulated differentially expressed genes (DEGs), while the lung samples had a smaller number of DEGs, mainly due to sample variability, so tissue-specific responses are likely “washed out” and only the generic responses (ISG, macrophages etc.) remain.

The tissues exhibited both shared and private gene sets, underscoring a coordinated yet locally-tuned innate program. The central observation of this study is the contrast between the strong immune response in peripheral tissues and the near-zero viral burden there: this is the transcriptional signature of successful clearance, not failed infection.

An alternative explanation could be that the response is cytokine and chemokine driven by the signal from the infected brain, which is plausible, but the combination of strong M2 signals, which are usually a sign of a restorative macrophage program after clearance of infection, along with strong induction of various primary innate response genes better supports our interpretation of a cleared viral infection.

### 4.4 Brain Immune Privilege, Not Neurotropism, Explains the Viral Distribution

IAV entry depends on sialic acid receptors synthesized by sialyltransferases: ST3GAL3/4/6 produce *α*2,3-linked (avian-type) receptors, ST6GAL1/2 produce *α*2,6-linked (human-type) receptors, and SLC35A1 transports sialic acid into the Golgi for glycoprotein attachment[39, 40, 41].

These genes are expressed uniformly across brain, kidney, and lung in uninfected embryos (Figure 4B), establishing that the virus has equal biochemical access to all three tissues. Despite this, viral transcripts at 48 h post-infection are overwhelmingly concentrated in brain and near-absent in kidney and lung compared to their own controls(Figure 4A). The conventional reading of this pattern is neurotropism, implying a preference of the virus for neural tissue. We argue the opposite: the uniform receptor data show the virus is not molecularly excluded from peripheral tissues. It has been cleared from them immunologically, and it persists in the brain because the brain is inaccessible to the circulating effectors responsible for that clearance. This reinterpretation rests on four converging lines of evidence. First, viral transcripts at 48 h postinfection are overwhelmingly concentrated in brain and near-absent in kidney and lung, indistinguishable from mock-infected controls, which suggests clearance, not low-level infection. Second, macrophage markers (CD163, CD206/MRC1,MMR1L4/MRC1L-B) are upregulated in peripheral tissues but conspicuously absent in brain, precisely the pattern expected if these effectors operate in the periphery but cannot enter the brain. Third, the macrophage populations in kidney and lung are M2-polarized which is the restorative phenotype that follows, rather than precedes, viral clearance. Fourth, as noted above, universal receptor expression(Figure 4) rules out differential receptor availability as an alternative explanation.

**Figure 4:**
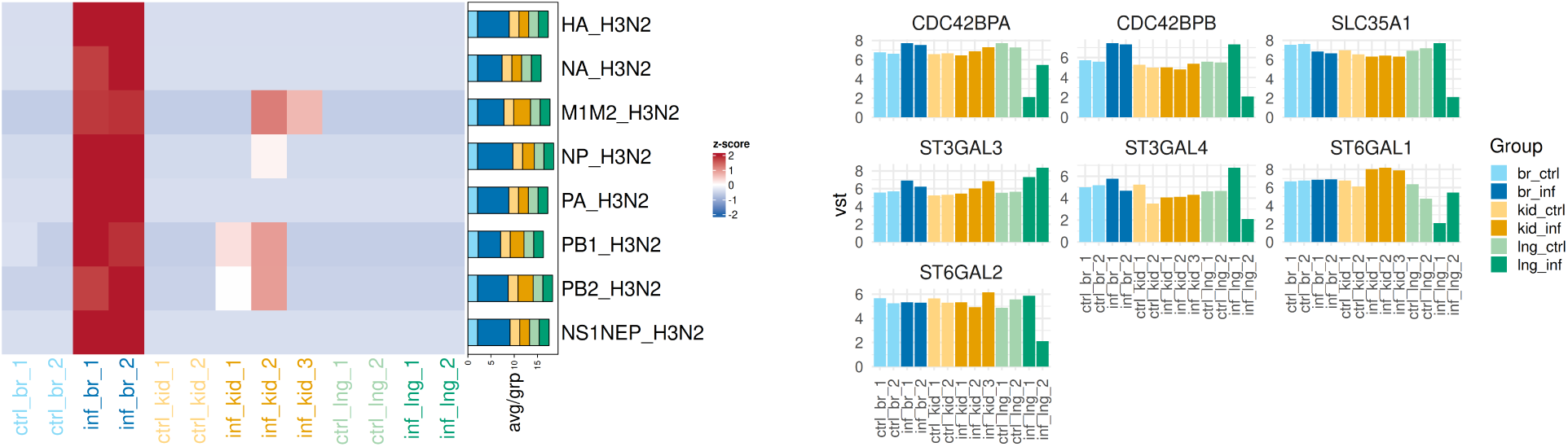
Viral concentration in the brain reflects immune privilege, not intrinsic neurotropism. **A)** Heatmap of normalized viral transcript counts for all 8 segments of the H3N2 genome across brain, kidney, and lung from infected and uninfected embryos. Viral transcripts are near-absent in infected kidney and lung at 48 h relative to their own uninfected controls, consistent with clearance. **B)** Expression of sialic acid related receptors that facilitate viral entry, including ST3GAL3, ST3GAL4, ST6GAL1, ST6GAL2, and SLC35A1 (H3N2 entry receptors) and CDC42BPA/CDC42BPB (which remodel the cytoskeleton) expression is present in all uninfected controls, ruling out differential receptor or cytoskeletal access as an explanation for brain-restricted viral burden[43, 44]. CDC42BPA and CDC42BPB are significantly upregulated in infected brain samples. ST3GAL3 is additionally upregulated in infected brain, potentially reflecting increased viral receptor production during active infection.

The logic here is analogous to the insight behind survivorship bias: during World War II, analysts initially concluded from studying bullet-hole patterns on returning aircraft that the marked areas, wings and fuselage, were the most vulnerable and should be reinforced. Abraham Wald recognized the error: the planes returning with holes in those areas were the survivors; the planes shot in the engines never returned at all. Viral absence in kidney and lung is similarly the mark of resilience, indicating those tissues cleared the infection. The brain, like the engine, is the site where the system failed, not because the virus preferred it, but because protection did not reach it.

Although full BBB maturation (dependent on endothelial tight junctions and basal lamina formation) is not complete until around E13[42]. Consistent with our detection of viral infection in the brain, macrophage markers (CD163, CD206/MRC1) that are robustly induced in kidney and lung are essentially absent from brain regardless of infection status, suggesting that the partial barrier structures present at E10 may be sufficient to exclude circulating macrophages from the brain parenchyma.

TLR2a, TLR4, TLR3, and avian-specific TLR21 are activated in the brain, indicating that the tissue mounts a meaningful innate response to infection despite the absence of circulating effectors. Neurodevelopmental transcription is maintained alongside immune activation: neuroblast differentiation genes (CCND1, PAX6, SIX3) remain elevated during infection, and CTNNB1 (*β*-catenin) induction suggests on-going Wnt-mediated neurogenesis(Figure SF5). This pathway intersects GSK3*β* co-elevation with antiviral signaling through phosphorylated *β*-catenin which promotes IRF3-dependent ISG expression (CXCL14, MX1, OASL)[45], which suggests there is simultaneous fine-tuned regulation. RTN4, an inhibitor of neurite outgrowth, is upregulated, potentially limiting neuronal communication as a damage-containment response. That the brain sustains both active neurogenesis and an antiviral program simultaneously underscores the embryo-protective character of this response, as well as its insufficiency for clearing the virus without immune cell infiltration.

### 4.5 Robust Antiviral Sensing via TLR–IRF Signaling and MDA5, Despite Absence of RIG-I

RIG-I (DDX58) is absent from the Galliformes genome[5, 6, 7], and its loss has been proposed as a molecular basis for heightened avian susceptibility to influenza. Our data refute this directly. Viral clearance in kidney and lung is complete at 48 h, and the downstream ISG response is fully activated (Figure 5), demonstrating that RIG-I is not required for effective peripheral clearance of H3N2 virus in this system.

**Figure 5:**
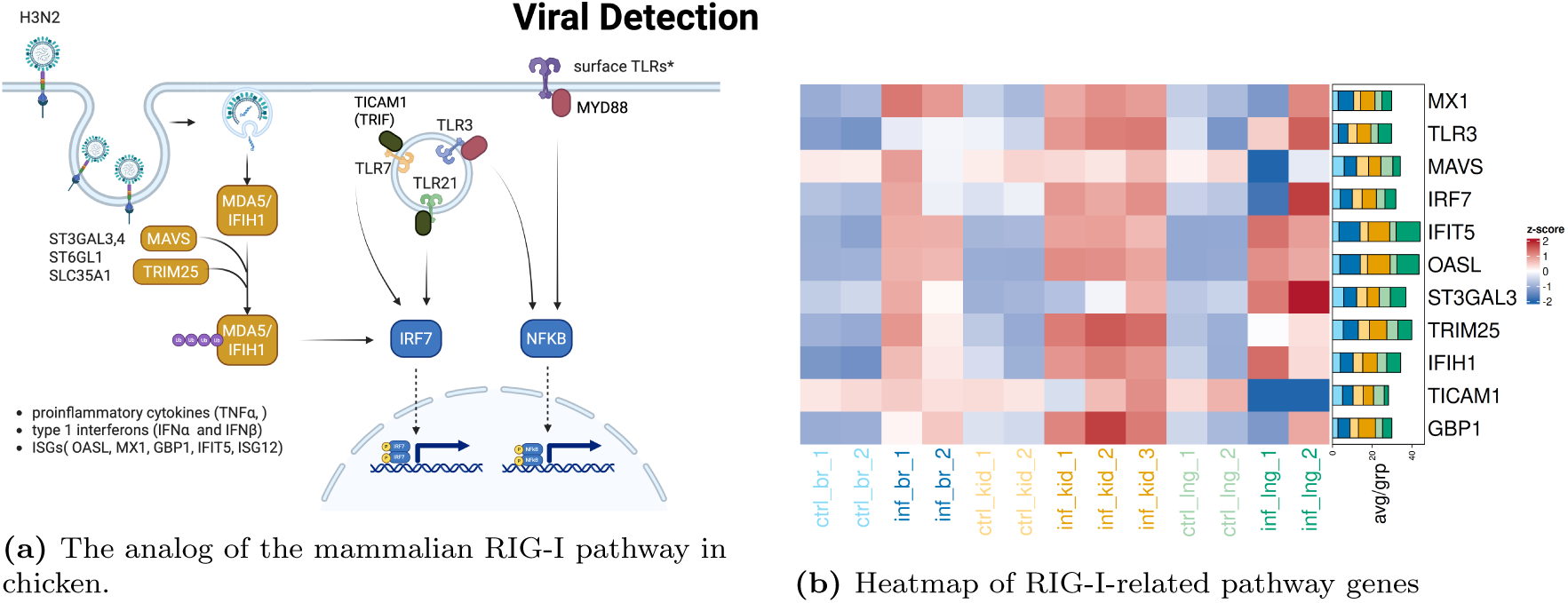
Functional antiviral sensing despite RIG-I absence. **a)** Influenza detection occurs though two main pathways: IFIH1/MDA5 and Toll-like receptor (TLR) signaling. Influenza invades and enters host cell through sialic acid receptors and its viral particles are released. IFIH1(MDA5) seemingly takes on the role of RIG-I and is activated upon recognition of viral RNA. Downstream factors, MAVS and TRIM25, are recruited and activate expression of pro-inflammatory cytokines via IRF7 signaling. In parallel, surface (TLR1, TLR2, TLR4, TLR5, TLR15, and TLR21) and endosomal (TLR3, TLR7, and TLR21) TLRs recognize influenza virion through each TLR’s complementary ligand, subsequently triggering IRF7 and NFkB pathways respectively. Cytokines from infected brain upregulate NAMPT (Nicotinamide phosphoribosyltransferase), which acts as a pivotal regulator of inflammation[48]. **b)** The heatmap shows most of the genes in panel A are upregulated in the infected brain and kidney samples, which in conjunction with data from Fig. 4 showing virus mostly in the brain consistent with the response in kidney being driven in part by systemic chemokine/cytokine signals from the infected brain rather than local viral infection.

Two parallel sensing arms compensate for RIG-I absence (Figure 5). First, MDA5 (IFIH1), which shares key structural domains and functional RNA-sensing activity with RIG-I, is highly expressed and serves as the likely primary cytosolic H3N2 sensor. TRIM25, which ubiquitinates RIG-I in mammals to potentiate downstream signaling, is elevated in both infected tissues; lacking its canonical substrate, TRIM25 here likely facilitates MDA5/IFIH1 activation instead[46, 47]. MAVS, IRF7, and downstream ISGs (OASL, MX1, GBP1, IFIT5, ISG12) are strongly induced, confirming a complete cytosolic sensing-to-effector axis. Second, and in parallel, the TLR-IRF axis provides endosomal RNA sensing. TLR3 (dsRNA) and TLR7 (ssRNA) are elevated in kidney, while TLR3 and the avian-specific TLR21 are activated in brain. TLR3 signals through TRIF (TICAM1) to activate IRF3; TLR7 and TLR21 signal through MyD88 to activate IRF7[13, 49]. IRF3 and IRF7 both regulate the type I interferon signaling pathway. Constitutive expression of IRF3 determines early stage viral infection and IFN beta production while IRF7 induced expression leads to interferon alpha production in mammals (Supplemental Figure SF2). IRF7 is a key early-response transcription factor in chicken macrophages, driving IFITM proteins and PKR/OAS family ISGs that directly inhibit viral replication. In our avian model, we see IRF7 being the main driver in IFIH1(MDA5) signaling and no significant expression of IRF3 in infected tissue(Figure SF2).

Despite the absence of RIG-I, we observe robust activation of downstream antiviral gene expression, consistent with functional compensation by MDA5 and other pattern-recognition pathways (chMDA5/chCARDIF/chLGP2 signaling), in line with prior transcriptomic evidence of MDA5-driven responses to HPAI strains H7N9, H7N1, H5N1, and H9N2[50, 51, 52, 53].

BF1 and BF2, the MHC class I genes central to avian antiviral defense, are highly upregulated in kidney but absent in brain[54]: BF1 presents antigens for Natural Killer (NK) cell surveillance and BF2 for Cytotoxic T Lymphocytes (CTLs), and their absence in the brain is consistent with its exclusion from immune cell patrolling. These MHC class I responses are therefore prospective rather than immediately functional at this developmental stage, positioned to support CTL and NK surveillance once adaptive competence matures. The full set of avian gene gains/losses and their functional compensators is summarized in Table ST2 and Table ST3; in most cases a functional substitute is present and active.

The kidney response provides additional evidence of systemic immune coordination driven by these sensing axes. Acute-phase proteins SAA1, ORM1, and S100A12 are strongly induced, alongside the biotin-sequestration proteins Avidin (AVD) and Biotinidase (BTD), which restrict pathogen access to biotin (Figure SF4, Table ST1). Notably, transferrin(TF) and albumin (ALB), which are classical negative acute-phase proteins normally suppressed during active inflammation, are paradoxically upregulated in infected kidney, a pattern consistent with a resolving rather than active infection. The chemokines CCL5, CXCL14, and CXCL12 are strong candidates for the systemic signals that relay information about brain infection to peripheral tissues across the blood-brain barrier. Kidney shows strong upregulation of NAMPT (Nicotinamide phosphoribosyltransferase), which is induced by pro-inflammatory cytokines (e.g. TNF-*α*, IL-1*β*, IL-6) from the infected brain and acts as a pivotal regulator of inflammation. Increased NAMPT, both intra- and extracellularly, fuels inflammation by increasing NAD+ for metabolic demands[48].

Iron management is a central thread connecting the sensing response to macrophage polarization (Figure 6). During active infection, iron is sequestered to starve replicating virus; free heme released from damaged cells is scavenged to prevent oxidative injury; and the transition from the M1 (inflammatory) to M2 (restorative) macrophage state is itself partly regulated by intracellular iron availability. All three processes leave transcriptional signatures in our data.

**Figure 6:**
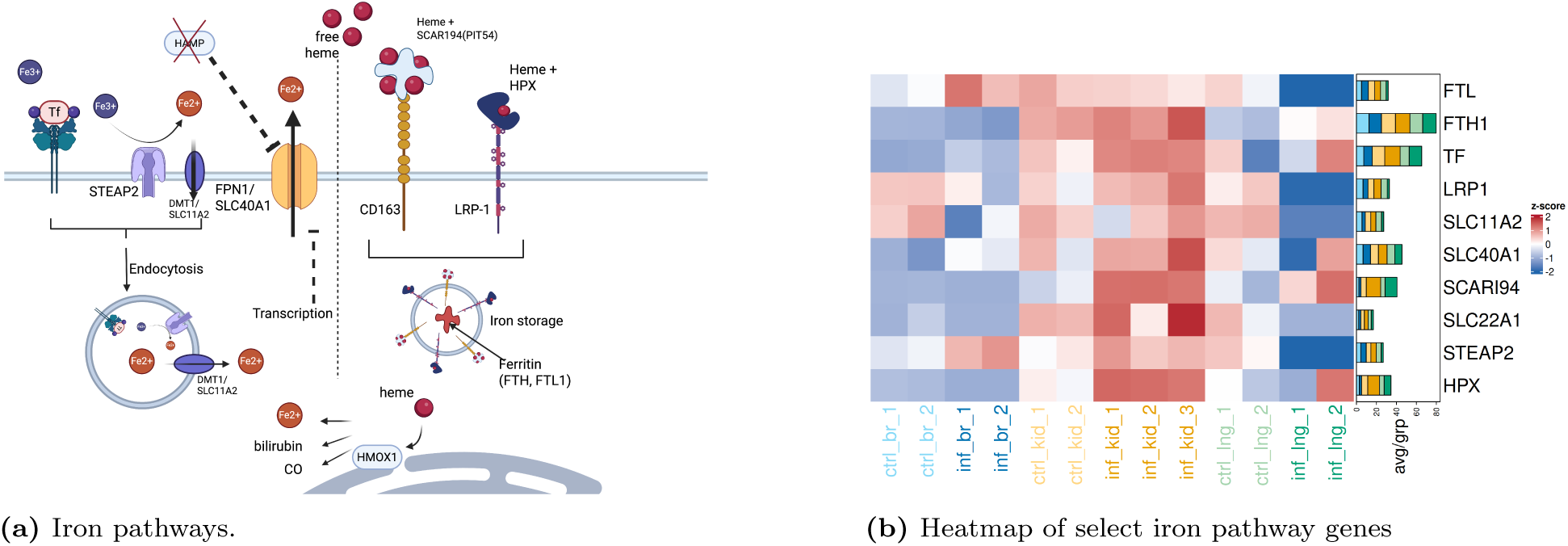
Proposed iron-sequestration pathway in chicken macrophages inferred from function of homologous proteins in mammals. **a)** Iron control in chicken inferred from a combination of mammalian pathways and chicken-specific studies (e.g. SCARI94). SLC40A1 (FPN1), the sole exporter of iron, is usually controlled by HAMP which is non-existent in chicken, suggesting the control could be transcriptional. Free heme outside the cell is captured by either SCARI94(the chicken equivalent of mammalian haptoglobin) or HPX(hemopexin) and transported into macrophages through either CD163(SCARI94 bound heme - hypothesized), or LRP1 (HPX-bound heme). Transferrin (TF) can also transport free iron in through endocytosis with the involvement of STEAP2 and SLC11A2 (DMT1). Ferritin (FTH1, FTL) plays a key role in sequestering iron in complexes for storage. Created in BioRender. **b)** Heatmap of iron-pathway genes. The data is consistent with macrophages being present in infected kidney but not in the infected brain. The up-regulation of TF, HPX, CD163, and the transporters, but not Ferritin suggest the urgent need to mop up heme is set in motion, but long-term storage might respond with a lag. SLC40A1 not being down-regulated suggests the control of iron as means of controlling infections might have some gaps compared to mammals, potentially allowing for viral replication.

### 4.6 Macrophages as the Primary Effectors of Clearance

The innate cellular response in peripheral tissues is dominated by restorative macrophage activity which is a phenotypic profile that characterizes resolved rather than ongoing infection (Figure 7).

**Figure 7:**
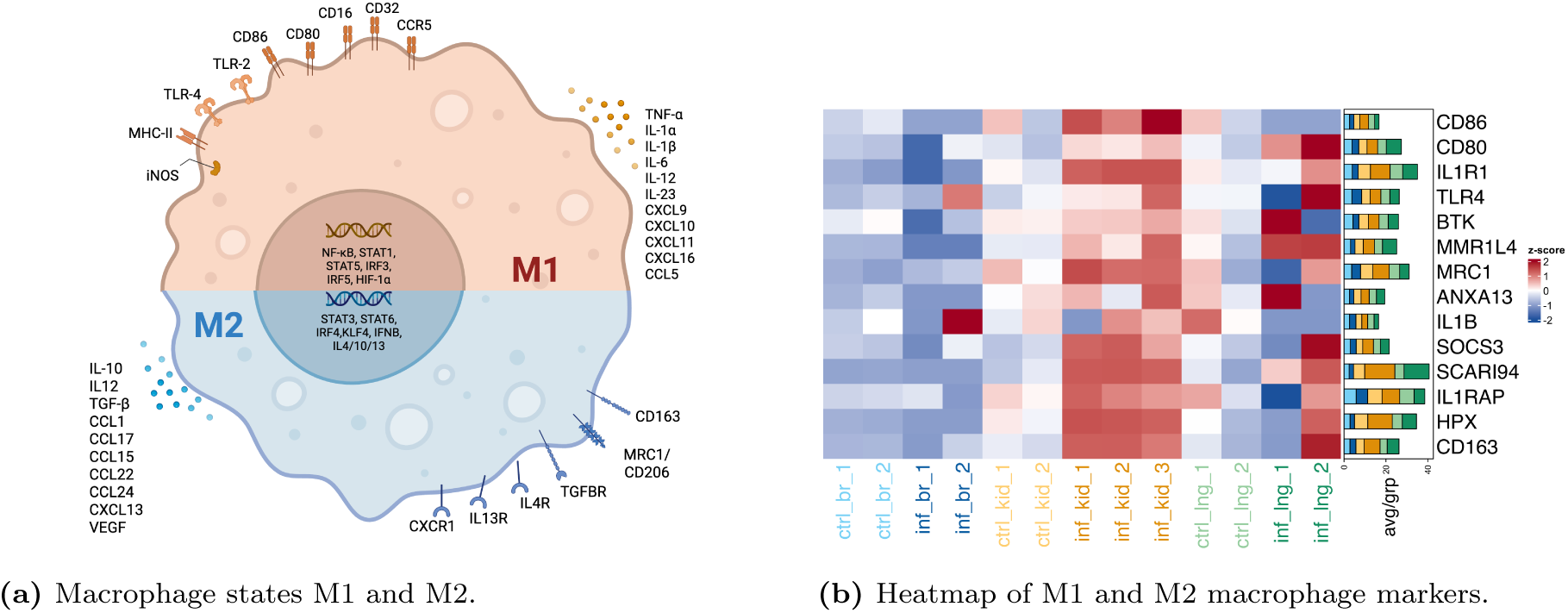
Macrophages as primary effectors. **a)** Macrophages can exist in two states/polarizations, M1 (brown) and M2 (blue), involving activation of different programs. Under infection, the M1 state helps clear infection, while the M2 states helps reconstruction and healing after clearance of the infection. Image created in Biorender. **b)** Heatmap shows M1 and M2 marker genes across tissues and conditions. In the infected kidney M1 genes (CD80, CD86) show very low expression, while exhibiting a marked induction of M2 markers (CD163, SCARI94, HPX, CD206/MRC1), which is the phenotypic profile of an infection that is being resolved. CD68, a common mammalian macrophage marker, is absent in chicken. Avian-specific MMR1L4 (MRC1L-B), which is a marker for macrophages, is present outside the brain and up in infected samples, suggesting recruitment of macrophages upon infection.

### M2 macrophage polarization

CD163, the hallmark M2/scavenger receptor, is elevated in kidney, while pro-inflammatory M1 markers (CD80, CD86) are largely absent. The functional equivalence of CD163 in chicken has not been directly demonstrated; we interpret its upregulation by analogy with mammalian M2 macrophage biology, which may not fully translate to the avian context. MRC1 (CD206) is upregulated in kidney and to a smaller extent in brain. Hemoglobin/heme detoxification is coordinated in the kidney by the strong upregulation of CD163, HPX, and SCARI94 (PIT54), the avian functional homolog of haptoglobin[55](Figure 6). ORM1 induction further links the acute-phase response to M2 polarization by promoting CD163 expression. The absence of M1 markers alongside active M2 markers is inconsistent with active viral infection requiring immediate inflammatory containment, instead, it is the signature of a post-clearance tissue-repair program (Figure 7).

HPX, released by macrophages as a second-line defense when haptoglobin (SCARI94) is depleted, scavenges free heme and delivers it via CD91/LRP1 internalization, preventing heme overload and suppressing proinflammatory macrophage activation[56].

Suppressors of cytokine signaling (SOCS) proteins regulate JAK/STAT pathways and macrophage polarization[57, 58]: SOCS1 suppresses JAK2/STAT1 in M1 macrophages, while SOCS3 inhibits JAK/STAT in M2 cells. Our study shows elevated expression of the SOCS family in both infected brain and kidney tissues. Elevated expression of SOCS1, SOCS2, and SOCS5 are dominant in infected brain.

SOCS3, elevated in infected kidney, may be driving M2 polarization via the CCL5/CCR5 axis[59, 60]. Despite elevated CCL5, its receptor CCR5 is downregulated, consistent with SOCS3 suppressing downstream JAK/STAT activation and steering macrophages toward inflammatory resolution.

### Complement is activated but lacks downstream efficacy

Classical complement components C1q, C1r, and C1s are strongly expressed in kidney, with the inhibitory C4BPA downregulated, indicating pathway engagement (Figure SF3). Lectin (MASP2) and alternative pathway (Factor I, Factor H) components are also elevated. However, the biological impact of this activation is likely limited. At day 10, embryos cannot synthesize antibodies de novo, and maternally transferred IgY has not been exposed to H3N2, which means no H3N2-specific antibody complexes are available to activate C1 via the classical route. C9, required for membrane attack complex (MAC) pore formation, also appears absent from the chicken genome. Complement activation here therefore probably represents a non-specific or pattern-based response rather than targeted viral opsonization, with no productive downstream lytic or antibody-dependent effector function.

### Microglia in the brain

In contrast to the peripheral tissues, the brain contains resident microglia from early developmental stages. Microglial markers (CD44, SPI1, CSF1R, PTPRC/CD45) are present in brain tissue, but circulating macrophage markers are absent. This confirms that the brain’s immune response relies solely on resident cells, and that the circulating M2 macrophages driving peripheral clearance are excluded from the CNS compartment, which is a defining feature of immune privilege.

Microglia are the primary resident phagocytes of the CNS and are expected to upregulate TREM2 upon infection[61, 62] which is not elevated in the infected brain samples relative to controls, in contrast to the strong activation signature (CD163,SCARI94 elevation) seen in peripheral macrophages; we return to this observation in the Discussion (Figure 8). CD68, a common mammalian macrophage marker, is absent in chicken. Avian-specific MMR1L4 (also known as MRC1L-B) is used as a macrophage marker and is present and responsive outside the brain, again suggesting exclusion of macrophages from the brain.

**Figure 8:**
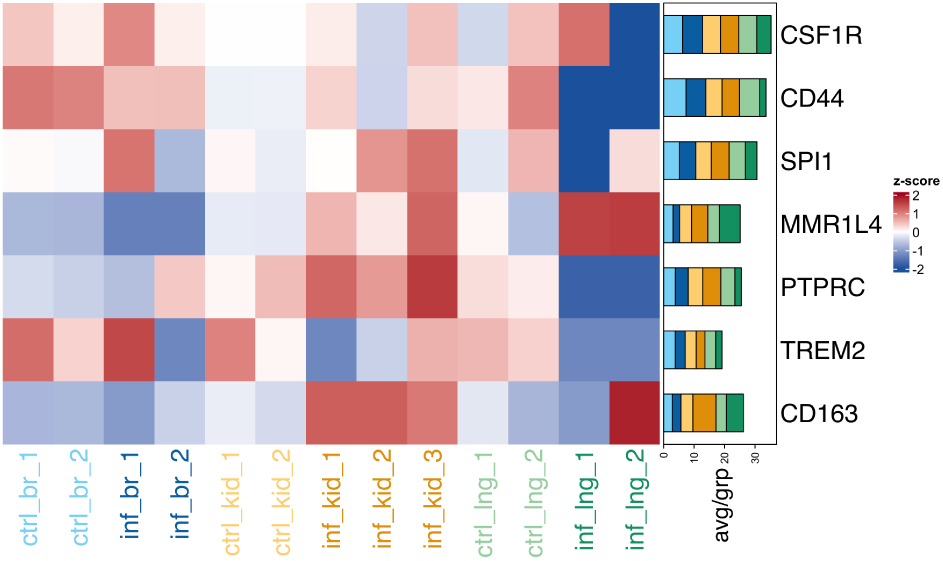
Microglia markers in brain. The microglia should mount a response to the infection, but apparently fail to do so in the 10-day old embryo. Microglial markers (CD44, SPI1, CSF1R, PTPRC/CD45) are present in the brain. TREM2, is expected to be induced upon microglial activation, show little or no change with infection. The circulating macrophage marker CD163 and chicken-specific MMR1L4 is responsive in infected kidneys but absent in brain, suggesting that circulating macrophages are excluded from the CNS and that the brain’s innate response relies on resident cells that fail to mount an activation response.

The microglia do not express several important markers such as SCARI94, MRC1, and MMR1L4 (the target of the antibody laboratory antibody called KUL01)[63], which are pattern-recognition receptors that bind high-mannose structures on viruses, bacteria, and fungi. These are upregulated primarily during the resolution or alternative activation (M2) phase rather than acute pro-inflammatory responses and are controlled by anti-inflammatory cytokines like IL-4 and IL-13.

### 4.7 Autophagy over Apoptosis as an Embryo-Protective Strategy

GBP1, an IFN*γ*-inducible effector, promotes autophagy and pathogen encapsulation while suppressing apoptosis; its strong upregulation across tissues is consistent with a system-wide preference for non-lytic pathogen clearance (Figure SF6). In the brain, ferroptotic signals are finely balanced: pro-ferroptotic indicators (GPX4 downregulation, ACSL4 elevation, TFRC upregulation) coexist with anti-ferroptotic compensators (elevated HIF1*α*, NCOA4)[64], suggesting regulated iron recycling via ferritinophagy rather than lytic cell death. NFE2L2 (NRF2)[65], metallothioneins (MT1/2/3), hemopexin (HPX), and STEAP family members are upregulated in brain, collectively limiting oxidative damage and constraining pathogen access to metals(Figure SF6).

In mammals, HAMP-mediated iron sequestration is a well-conserved component of nutritional immunity[66, 67, 68]. The absence of this axis in the chicken embryo is a potential vulnerability: elevated bioavailable iron in the brain may contribute to the unrestrained viral infection observed there. Ceruloplasmin (CP), a ferroxidase, is highly upregulated in kidney and acts to reduce tissue damage and mobilize iron away from pathogens; its absence in brain represents a further gap in the local metal-withholding defense. Whether the HAMP deficiency represents a transient developmental feature or a fixed aspect of avian physiology warrants direct investigation.

## 5 Discussion

### 1. The embryonic innate immune system is likely sufficient for viral clearance at least for chicken-adapted H3N2

The central finding of this study is that a 10-day chicken embryo, which lacks mature T cells, B cells, and any adaptive immune capacity, can clear systemic H3N2 influenza infection through innate mechanisms alone. This is demonstrated directly by the near-zero viral burden in kidney and lung at 48 h, in tissues that simultaneously show robust transcriptional immune activation. This is not the profile of a host that failed to respond; it is the profile of one that responded and succeeded. The innate toolkit consisting of MDA5/IFIH1, TLR3-TLR7-IRF signaling, and M2-polarized macrophage is sufficient to control and clear a human influenza strain in the absence of any adaptive contribution. Complement is activated but, as discussed below, lacks the H3N2-specific antibodies needed for productive downstream function. The full PRR landscape in poultry is summarized in Supplementary Figure SF2.

### 2. RIG-I absence does not impair innate antiviral defense against H3N2

The hypothesis that Galliformes are more susceptible to influenza because they lack RIG-I[5, 6, 7] is called to question by our data. Peripheral clearance of H3N2 is complete, and the downstream ISG response is fully and robustly activated. MDA5/IFIH1 with TRIM25 co-activation reconstitutes the functional equivalent of the mammalian RIG-I–TRIM25 axis, while TLR3–TRIF and TLR7/TLR21–MyD88–IRF7 provide parallel and redundant sensing routes[13, 49]. Each avian-specific gene loss (eġİSG15, IFIT1/2, haptoglobin) is similarly paired with a functional avian substitute (OASL, IFIT5, SCARI94/PIT54). The chicken innate immune system has seemingly not passively lost RIG-I; it has likely evolved an alternative architecture that demonstrably achieves the same functional outcome, at least for H3N2. That reinstating RIG-I in chickens increases inflammation rather than protection[19] further argues that the existing avian architecture is not merely compensatory but may represent an optimized alternative. Studies attributing avian susceptibility to RIG-I absence might want to account for this evidence.

It is possible that the lack of RIG-I might render the chicken susceptible to certain viruses, but the simple picture of this leaving a gaping hole in the innate defence is called to question by our data.

### 3. Brain immune privilege, not neurotropism, explains the viral distribution

The concentration of virus in the brain does not reflect a preference of H3N2 for neural tissue. Rather, it likely reflects the fact that the brain is the one tissue in the embryo that circulating innate effectors cannot reach. The partial barrier structures at E10 are sufficient to exclude circulating macrophages from the brain parenchyma[42], and this is directly confirmed by our data: microglial markers are present in brain while circulating macrophage markers are absent. Universal expression of ST3GAL3, ST3GAL4, ST6GAL1, ST6GAL2, and, SLC35A1 across tissues confirms that viral entry is biochemically possible everywhere; the brain-restricted viral burden is therefore a product of immune architecture, not viral preference. Notably, the virus appears to be making no active effort to evade immunity as there is no evidence of ISG suppression or immune evasion gene expression. It suggests infection is possible when macrophages cannot reach the tissue. This has potential clinical relevance: the same logic may explain why influenza neurological complications are disproportionately reported in young children[69, 70, 71], whose peripheral innate systems are robust but whose developing brain-immune barrier may present a similar sanctuary.

There is a possibility that intrinsic tissue permissiveness to replication, which can depend on factors like functional receptor density, glycan branching/sialylation patterns, membrane fluidity, or local temperature, might explain viral presence in the brain, independent of any immune process. This could be addressed by time-series measurements that could show virus transiently infecting other tissues before being cleared.

### 4. Microglia in the chicken embryo brain do not respond to the infection

Microglia are the primary resident phagocytes of the CNS and, upon infection, would be expected to upregulate TREM2 as they shift from routine developmental clearance toward active pathogen engulfment[61, 62]. We don’t observe a response: TREM2 expression in infected brain does not differ appreciably from uninfected controls, even as the brain accumulates the highest viral burden of any tissue profiled. It is likely that the microglia, being yolk-sac/embryonically derived rather than monocyte-derived, may not yet be primed to mount an M2-like or phagocytic response at this developmental stage. We offer this as a plausible, cell-intrinsic explanation for brain viral persistence that complements the tissue-access argument above; it remains inferential from transcriptomic markers, and direct validation (e.g., microglial immunostaining or functional phagocytosis assays across developmental stages) would be needed to establish it.

### 5. A restorative program preserves developmental integrity

The dominance of M2 macrophage polarization and autophagy over apoptosis reflects a host strategy that resolves infection while minimizing collateral developmental damage. The atypical upregulation of negative acute-phase proteins (TF, ALB) in kidney further supports the interpretation that the peripheral infection is resolving rather than progressing. The collapse of HAMP-mediated iron withholding[66] remains a potential vulnerability, and its contribution to unrestrained brain infection (and possible replication) warrants direct experimental investigation. That ceruloplasmin compensates in kidney but is absent in brain suggests that the iron-management gap is specifically a CNS vulnerability, compounding the immune exclusion imposed by the blood-brain barrier (Figure SF6).

### 5.1 Limitations and Future Directions

The small number of samples is a big limitation of our study. Repeating this with many more samples would be an important follow-up. A time-series after infection, over the few hours leading upto the 48-hour time point of our study, would help delineate the course of infection clearance in kidneys and develop a better picture of the response. *Orthogonal validation by staining, flow cytometry (immunostaining or flow cytometry for immune-cell infiltration, and functional pathway perturbation)* would remove any ambiguity regarding tissue-specificity of infection. Another useful study would be to infect the mother with the same virus and study the response of the 10-day embryonated eggs to the infection, to see the role the complement plays in viral defense. Using a wild-type avian influenza virus instead of the adapted human virus we used would also help strengthen the conclusions. Our study is a first step, providing landmarks that will guide future studies that are bound to follow.

## 6 Conclusion

Embryonated chicken eggs mount an innate immune response to H3N2 influenza that is sufficient to clear systemic infection without any adaptive contribution and they achieve this despite lacking RIG-I, a sensor previously invoked as central to avian influenza susceptibility. Two parallel sensing arms, consisting of cytosolic MDA5/IFIH1 and endosomal TLR3/TLR7–IRF signaling, drive a robust ISG response, and M2-polarized macrophages execute clearance in peripheral tissues. Complement is activated but cannot contribute productively: the embryo has no H3N2-specific antibodies and lacks C9 for membrane attack complex formation. Viral persistence is therefore confined to the brain, an immune-privileged sanctuary where macrophages cannot penetrate, consistent with resident microglia that appear not yet primed to mount a response at this developmental stage, and the conspicuous absence of infiltrating macrophage markers at the primary infection site. These findings suggest a reframing of the significance of avian RIG-I loss, indicate the embryonic brain is likely an early-life immune-privileged compartment, and illuminate a previously uncharacterized innate immune architecture in a host of major agricultural and epidemiological importance.

## Acknowledgements

Asif Khalak provided early criticism and suggestions that were helpful. The three anonymous reviewers at *Review Commons* were exceptionally helpful with their criticisms, suggestions, and encouragements.

## Glossary

APP: Acute phase proteins
BBB: Blood-brain barrier
CP: Ceruloplasmin; a copper-binding ferroxidase involved in iron oxidation and transport
CTL: Cytotoxic T Lymphocytes
DEG: Differentially expressed gene
ECE: Embryonated chicken egg
EID_50_: Egg Infective Dose that infects 50% of eggs in a test group
HA: Hemagglutinin
HA assay: Hemagglutination assay
IFN: Interferon
IRF: Interferon regulatory factor
ISG: Interferon-stimulated gene
MAC: Membrane attack complex
MDA5: Melanoma differentiation-associated protein 5; officially IFIH1. Primary cytosolic RNA sensor in chickens, compensating for absent RIG-I
NAMPT: Nicotinamide phosphoribosyltransferase
NK: Natural Killer
PAMP: Pathogen-Associated Molecular Patterns
PRR: Pattern recognition receptor
RIG-I: DDX58;Retinoic acid-inducible gene I; recognizes viral RNA and initiates the innate antiviral response; absent in Galliformes
RLR: RIG-I-like receptor
SPF: Specific pathogen-free
TLR: Toll-like receptor

## 7 Data Availability

Gene expression data and an interactive exploration tool are available at https://katahdin.girihlet.com/chicken_egg/

## S1 Supplementary Material

### S1.1 Acute Phase proteins in the chicken

Acute phase proteins (APPs), crucial components of innate immunity, are blood proteins (primarily produced in the liver), whose serum concentrations change by *≥* 25% during inflammation, infection, or tissue injury. Their transcription is driven by cytokines (IL-6, IL-1, TNF-*α*) and they function to destroy pathogens, activate complement, and promote repair. The chicken-specific APPs relevant to our study are shown in Table ST1.

**Table ST1:**
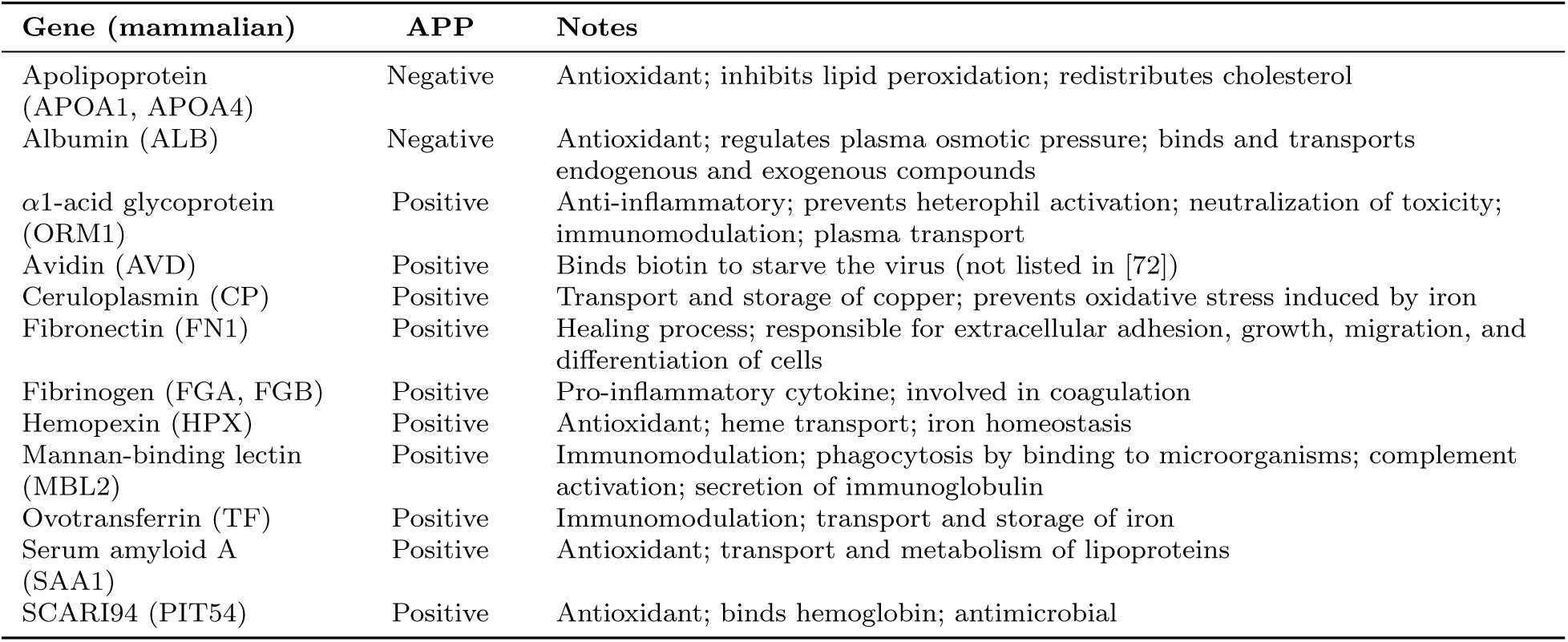
Acute phase proteins (APPs) from [72, 73].

### S1.2 Avian-specific immune gene losses and functional compensators

Several canonical mammalian immune genes are absent from the chicken genome. The losses most relevant to antiviral defense, and their avian functional replacements, are summarized in Table ST2. In each case a functional substitute is present and transcriptionally active, consistent with the complete peripheral viral clearance observed in this study.

Collectins (collagen-containing C-type lectins) are soluble pattern-recognition molecules of the innate immune system that identify and neutralize pathogens. They act as crucial first-line defenders by binding sugar patterns on microbes to promote aggregation, opsonization, complement activation, and inflammation modulation. There are significant differences between avian and mammalian collectins delineated in Table ST3.

### S1.3 Wnt/***β***-catenin and Neurodevelopmental Context

The Wnt/*β*-catenin pathway is a critical regulator of neuronal development, feather morphogenesis, and cell communication in chicken embryogenesis[74]. In infected brain, co-transcriptional activators cTCF and IRF3 are elevated; although LEF1 remains unchanged, cTCF may transcriptionally activate Wnt target genes. Concurrent elevation of GSK3*β* (which phosphorylates *β*-catenin for proteasomal degradation) suggests tight negative regulation despite activator induction. WFIKKN2 (a TGF*β* family transporter involved in Wnt/*β*-catenin signaling) was significantly enriched in infected brain. Independently, phosphorylated *β*-catenin can partner with IRF3 and p300/CBP to activate ISGs (CXCL14, MX1, OASL)[45], providing a mechanistic link between ongoing neurogenesis and antiviral transcription in the immune-privileged brain.

At day 10–11, neural tube formation is complete and the brain has distinct prosencephalon, mesencephalon, and rhombencephalon regions[75]. Neuroblast differentiation markers (CCND1, PAX6, SIX3) remain elevated during infection, while RTN4 (inhibitor of neurite outgrowth) is upregulated, potentially limiting neuronal communication as a damage-containment response. The maintenance of neurodevelopmental programs during active viral infection underscores the embryo-protective character of the brain’s intrinsic response, while also illustrating that this intrinsic response alone cannot clear the virus.

**Table ST2:** Avian-specific immune gene losses and functional compensators.

| Gene (mammalian) | In chicken | Compensator | In mammals | Notes |
| --- | --- | --- | --- | --- |
| RIG-I (DDX58) | **NO** | IFIH1 (MDA5) | YES | Lost in Galliformes; MDA5 shares structural/functional domains; downstream ISG response fully intact |
| ISG15 | **NO** | OASL | YES | OASL retains antiviral ISGylation-like activity |
| IFIT1/IFIT2 | **NO** | IFIT5 | YES | chIFIT5 binds 5'-triphosphate viral RNA |
| Haptoglobin | **NO** | PIT54 (SCARI94) | YES | Binds hemoglobin; preserves NO signaling |
| TLR9 | **NO** | TLR21 | YES | TLR21 recognizes CpG DNA; functional replacement confirmed |
| MBL2 | **YES** | — | Pseudogene | Chicken lung lectin; functional collectin |
| MX2 | **NO** | — | YES | Role in avian antiviral response unclear |
| Avidin (AVD) | **YES** | — | **NO** | Avian-specific; sequesters biotin to restrict pathogen growth |
| PIT54 (SCARI94) | **YES** | — | **NO** | 22 novel avian SCAR genes absent in mammals |
| C9 (MAC) | **NO?** | Unknown | YES | Absence may limit MAC lysis; complement likely functions in opsonization |
| HAMP (hepcidin) | **NO** | — | YES | Iron-withholding axis largely non-functional in embryo; potential CNS vulnerability |

**Table ST3:** Avian collectin genes and their mammalian counterparts. MBL2 was the most transcriptionally reactive collectin in infected embryos; others showed minimal induction, possibly reflecting embryonic immaturity.

| Avian name | Gene symbol | Notes |
| --- | --- | --- |
| Chicken MBL | *MBL1* | Primary lectin pathway initiator in birds; no response observed |
| Chicken Lung Lectin (cLL) | *MBL2* | Functional in chickens; humans carry a pseudogene; most reactive collectin |
| SP-A | *SFTPA1/2* | Surfactant protein A; minimal response |
| cCL-1 | *COLEC10* | Homolog of mammalian Collectin liver 1 |
| cCL-2 | *COLEC11* | Homolog of mammalian Collectin kidney 1 |
| cCL-3 | *COLEC12* | Homolog of mammalian CL-P1 |

### S1.4 Ferroptosis Signaling Detail

Ferroptosis in the infected brain shows competing signals (Figure 6). Upregulation of TRIM25 suppresses ferroptosis by activating NRF2 allowing cells to survive[76]. Other anti-ferroptotic signals include elevated HIF1*α* and NCOA4. PEBP1/RKIP1 (which interacts with LOX enzymes) is downregulated; CPEB4 is upregulated. Pro-ferroptotic indicators include: KEAP1 elevation (normally mediates NRF2 degradation), GPX4 downregulation (loss of key ferroptosis inhibitor), ACSL4 and SP1 elevation (fatty acid metabolism promoting ferroptosis), and TFRC elevation (iron importer, consistent with extrinsic pathway activation)[64]. We interpret the likely outcome as regulated iron recycling via ferritinophagy rather than lytic ferroptotic cell death, consistent with the broader autophagic program.

**Figure SF1:**
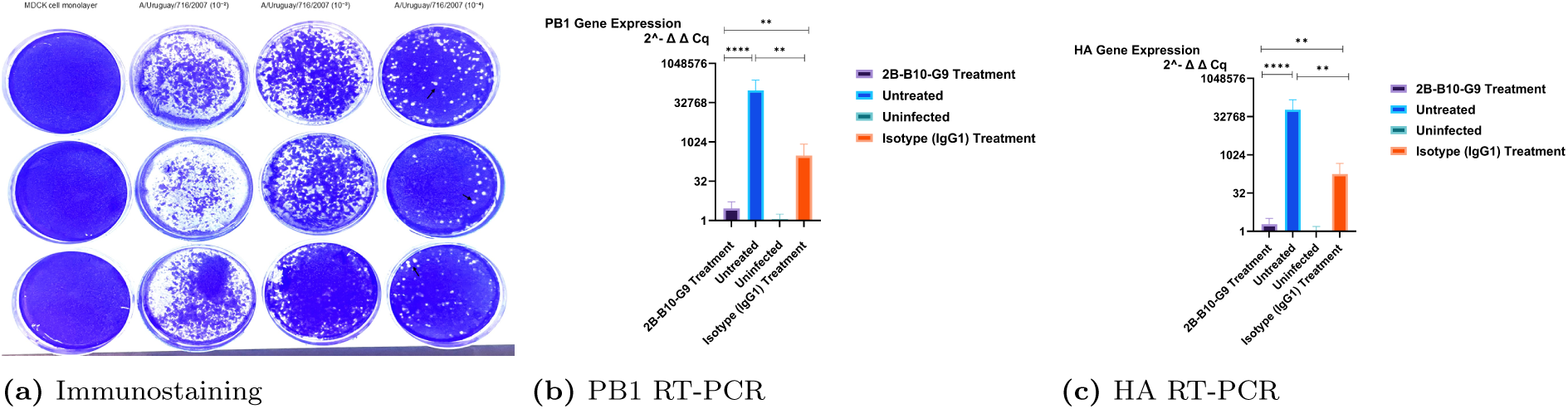
Confirmation of viral infection by immunostaining and RT-PCR. **Panel a:** In vitro replication and spread of virus to neighbouring MDCK-SIAT1 cells at different dilutions after the intact cells stained with crystal violet. This identifies the optimum dilution of virus for in ovo inoculation that is 60-65 countable distinct border plaques (each plaque is considered to be formed by one virus particle) after the plaques (white/unstained areas showing lysed/infected cells) were counted. **Panels (b,c):** In ovo replication of the virus in the epithelial cells of chorioallantoic membrane is shown by the HA and PB1 (polymerase basic 1- main polymerase of the virus for replication) gene expressions quantified by RT-qPCR of the allantoic fluid harvested after 48 hours of in ovo virus inoculation. The treated group with anti-matrix1 (M1) mAb showed reduction of virus replication via reduction of the HA and PB1 gene expressions in comparison to isotype IgG1 treated group. Since the antibody blocks replication, this result suggests there is active replication of the virus in the infected cells.

**Figure SF2:**
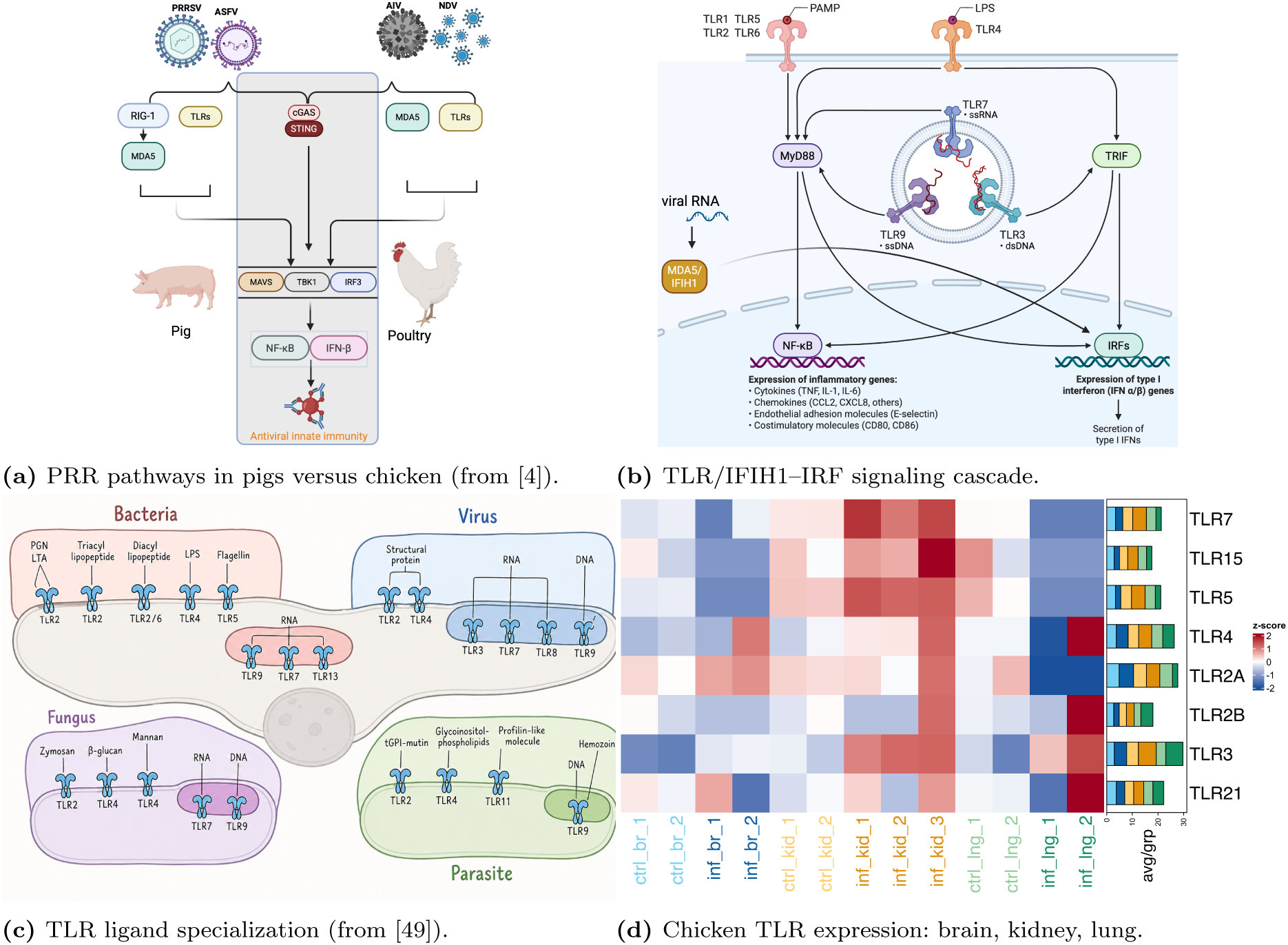
Innate sensing architecture in the chicken embryo. These eventually induce transcription of chemokines, cytokines and other interferon responsive genes that transmit signals to neighboring cells, alerting them to infections. **a)** Pattern recognition receptors (PRR) in pigs (mammals) versus chicken (Galliformes) include endosomal Toll-like receptors (TLRs), cytosolic RIG-I-like receptors (RLRs), e.g. MDA5, but not RIG-I in Galliformes. The PRR also include the cGAS–STING axis[4]. Many livestock viruses actively evade these pathways-AIV NS1 impairs RNA sensing, NDV V protein targets MDA5, and ASFV MGF proteins disrupt cGAS–STING-but H3N2 shows no such evasion activity here, consistent with a virus adapted to its human host. Created in BioRender. **b)** Interferon regulatory factors (IRF) are activated downstream, following TLR or IFIH1/MDA5 engagement, driving ISG transcription and the antiviral effector response. Created in BioRender. **c)** Toll-like receptor (TLR) ligand specialization across the receptor family, illustrating the range of pathogen-associated patterns detected[49]. Chicken TLRs include the avian-specific TLR15 and TLR21, the latter replacing mammalian TLR9 for CpG DNA detection[77]. **d)** Heatmap of chicken TLR expression across brain, kidney, and lung in infected and uninfected embryos. TLR2a, TLR4, TLR3, and TLR21 are activated in infected brain; TLR3, TLR5, TLR7, and TLR15 are elevated in infected kidney. The divergence between tissues likely reflects both their distinct TLR repertoires and their different infection states — active versus cleared infection.

**Figure SF3:**
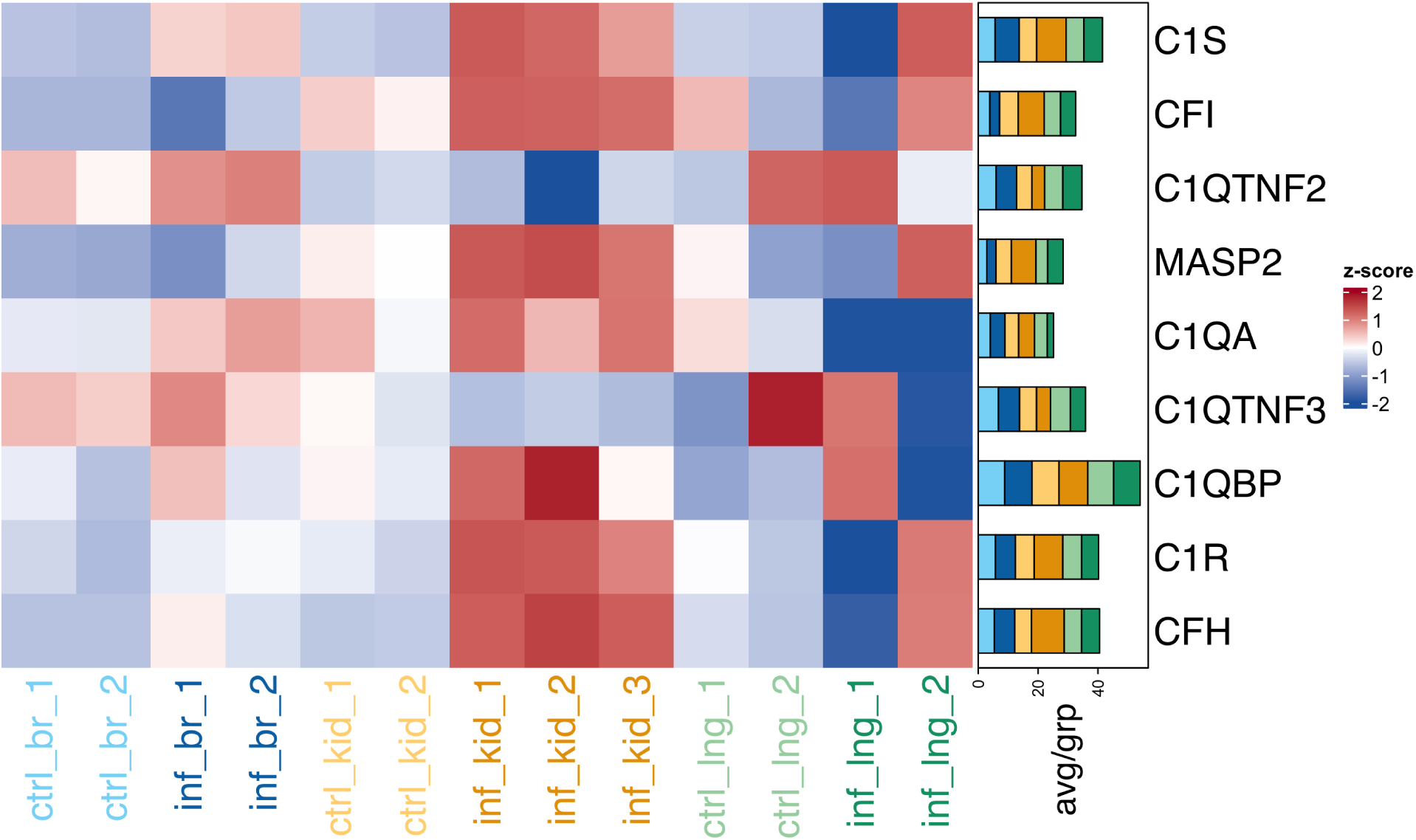
Complement pathway activation without productive downstream effect. Heatmap of complement pathway components (C1q, C1r, C1s, MASP2, Factor H, Factor I) in infected vs. uninfected kidney and brain, which represent the three complement pathways (classical/lectin/alternative), indicating multi-pathway engagement, though productive opsonization is likely limited by the absence of H3N2-specific antibodies.

**Figure SF4:**
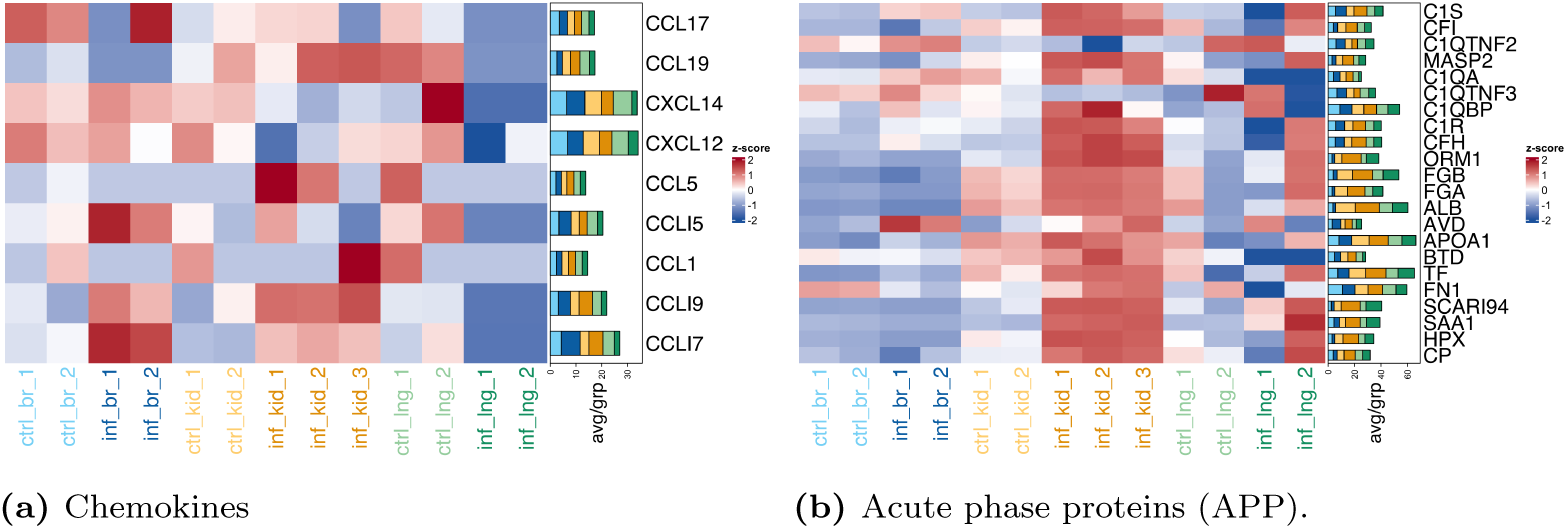
Chemokine and APP responses. **a)** Candidate chemokines (CCL5, CXCL14, CXCL12, CCL17, CCL19, CCL1) as potential mediators of systemic immune coordination from infected brain to peripheral tissues. CCL17 and CCL19 are upregulated in infected brain and kidney, while CCL15 and CXCL14 are only upregulated in the infected brain and the rest don’t seem to respond. **b)** Heatmap of acute-phase response (APP) genes (SAA1, ORM1, S100A12, TF, ALB) and biotin-sequestration proteins (AVD, BTD) across tissues and conditions[72, 73]. All the APP are upregulated in the infected kidney, but only AVD, TF, and APOA1 are up in the infected brain. Upregulation of negative acute-phase proteins (TF, ALB) in infected kidney is consistent with a resolving infection; AVD sequesters biotin to restrict pathogen growth.

**Figure SF5:**
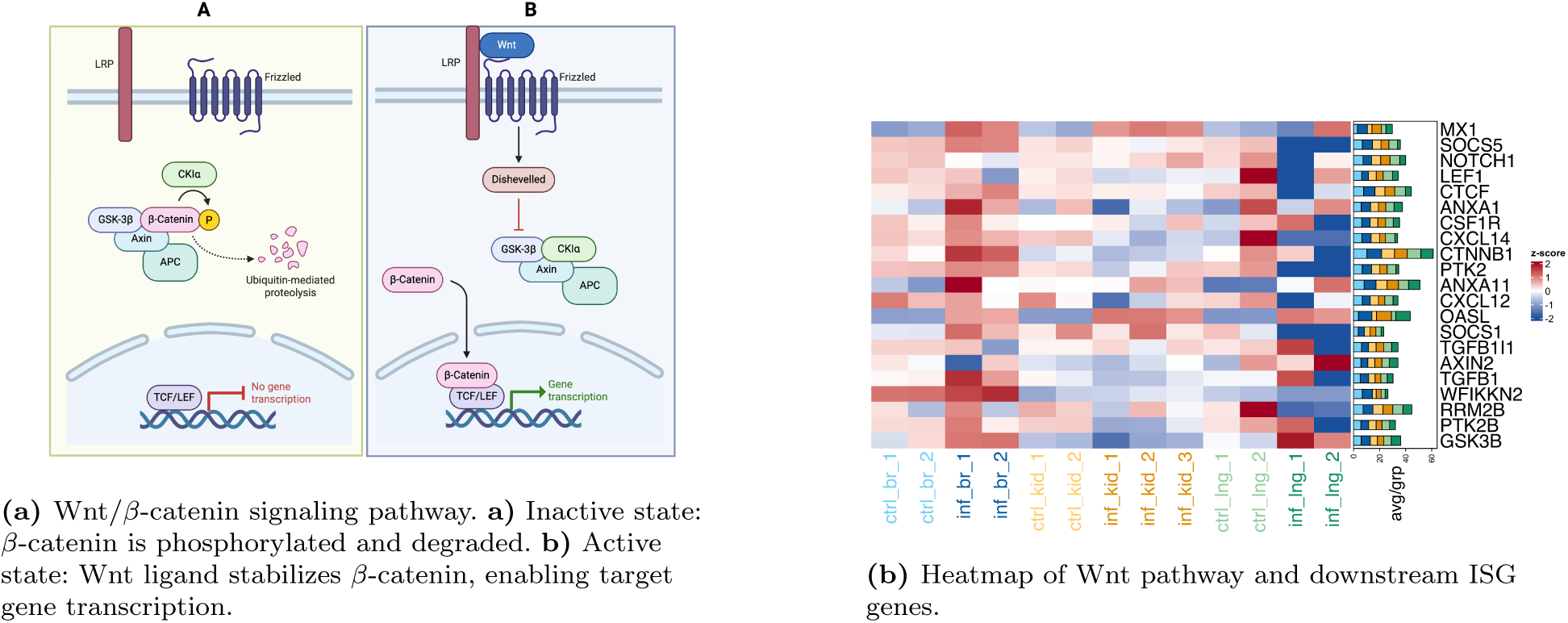
Wnt/*β*-catenin pathway gene expression in infected brain. a) Wnt signaling regulates critical developmental processes, such as neural patterning, organogenesis, and cell migration. The canonical pathway, in the *On* state, *β*-catenin gets stabilized and enters the nucleus to activate Wnt-target genes. Adapted from [78], created in BioRender. **b)** Heatmap of Wnt pathway components (CTNNB1, GSK3*β*, CTCF, LEF1, CCND1, AXIN2, WFIKKN2) and IRF3-linked ISGs activated downstream of *β*-catenin (CXCL14, MX1, OASL) in infected vs. uninfected brain. Most are upregulated in the infected brain, the concurrent elevation of activators and negative regulators reflects fine-tuned balance between ongoing neurogenesis and antiviral signaling in the immune-privileged brain. Note: CXCL10 is absent from the chicken genome; CXCL14 serves the analogous chemokine function here.

**Figure SF6:**
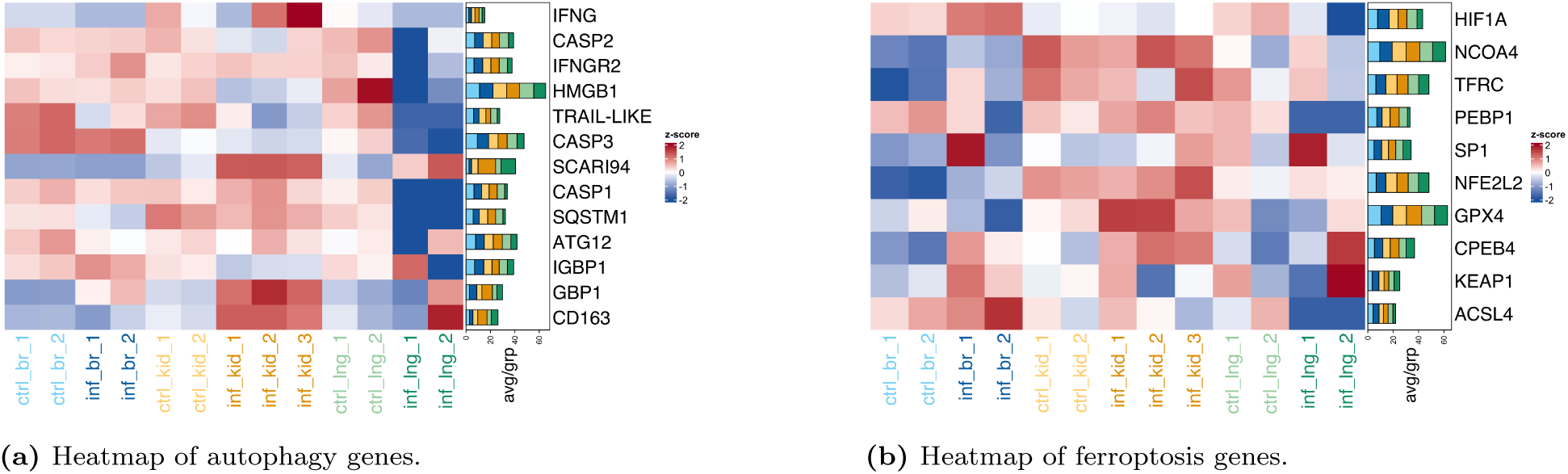
Preferential autophagy over apoptosis, with balanced ferroptotic signals in infected brain. **a)** Heatmap of autophagy. GBP1, SCARI94, CD163 are upregulated in infected kidney, while some others are up regulated in the infected brain (HMGB1, IGBP1) while others do not respond (SQSTM1,ATG family). There is no apoptotic response (TRAIL, caspases). Strong autophagy induction with suppressed apoptotic markers is consistent with a non-lytic embryo-protective clearance strategy. **b)** Heatmap of ferroptosis pathway genes (NCOA4, NFE2L2, TFRC, GPX4, PEBP1, ACSL4, HIF1*α*, SP1, KEAP1, CPEB4) across tissues and conditions. Competing pro-ferroptotic (GPX4 downregulation, ACSL4 and TFRC elevation) and anti-ferroptotic (HIF1*α*, NCOA4 elevation) signals in infected brain indicate regulated ferritinophagy rather than lytic cell death.

## Notes

https://katahdin.girihlet.com/chicken_egg/

### Competing Interest Statement

The authors have declared no competing interest.

### Summary of Updates

The list of substantial changes made in the manuscript includes; Revamped Figure 1 to remove superfluous panels, discussion of hemagglutination assay (HA assay) and edited the caption to more accurately reflect the work done. The Materials and Methods section has been thoroughly revised with new sections and more explanations; Improved discussion of virus used in the study. Added details for: Justifications for use of H3N2 virus strain in this study L194P mutation in the strain that allows infection of chicken HA assay and its use in our study Prior studies that detail methods used here Experiments to determine dosage Infection protocol Expanded details for the collection of genomic information (gene sequences and names) used in our study Detailed the mRNAseq analysis We added tissue-identity marker-gene validation (PAX6/SIX3/STMN3 for brain, SULT/FGG/FGB for kidney) to address the possibility of bad dissections at E10. We added data from plaque-assay/RT-qPCR-of-allantoic-fluid experiments. The corrected/replaced citation of the old H5 HPAIV review that didn't support our claim. Improved the legibility of all figures (heatmaps/plots) by increasing fonts and removing margins Better justified our claims of infection clearance in kidney/lungs using viral sequences detected in our samples. Toned down discussion to indicate this study is not exhaustive, but is indicative of a roadmap for further investigations. Provided more context to the RIG-I claims, citing alternate explanations for our findings. Reworded the abstract to emphasize that if even one virus, H3N2, is cleared by the innate immune system of the embryo without needing RIG-I, then the centrality of RIG-I as an essential component is called into question. Expanded discussion of role of microglia in the brain and their response to the infection in embryonated eggs The web-based tool has been redesigned from the ground up with several enhancements and new features allowing better exploration Added a download link for raw data and underlying genomic resources developed for this analysis Moved microglia heatmap to main text and added more markers and information. Added a figure to show tissue-specific expression as well as PCA plot to demonstrate integrity of kidney and brain dissections. Added a concluding section on limitations and future directions in the discussion

